# The evolution of life history theory: Bibliometric analysis of an interdisciplinary research area

**DOI:** 10.1101/510826

**Authors:** Daniel Nettle, Willem E. Frankenhuis

## Abstract

Life history theory developed as a branch of formal evolutionary theory concerned with the fitness consequences of allocating energy to reproduction, growth and self-maintenance across the life course. More recently, researchers have advocated its relevance to many psychological and social-science questions. As a scientific paradigm expands its range, its parts can become conceptually isolated from one another, so that in the end it is no longer held together by a common core of shared ideas. Here, we investigate the life history theory literature using quantitative bibliometric methods based on patterns of citation. We found that the literature up to and including 2010 was relatively coherent: it drew on a shared body of core references, and had only weak cluster divisions running along taxonomic lines. The post-2010 literature is more fragmented: it has more marked cluster boundaries, including boundaries within the literature on humans. Specifically, there are two clusters of human literature based around the idea of a fast-slow continuum of individual differences in behaviour that are bibliometrically isolated from the rest of the literature. We also find some evidence suggesting a relative decline in formal mathematical modelling. We point out that the human fast-slow continuum literature is conceptually closer to the non-human ‘pace of life’ literature than to the non-human literature usually referred to as ‘life history theory’.

## Introduction

Life history theory was developed in the mid to late 20^th^ century as a component of evolutionary theory. It offered tools for explaining variation in how organisms allocate energy to reproduction— or alternatively to other activities such as growth or somatic maintenance—across the life course, and across offspring. The development and testing of life history theory came to be a recognisable research programme in ecology and evolutionary biology. Subsequently, life history theory also began to be appealed to as a resource for answering questions in a range of human-focussed disciplines, such as psychology [1–4], anthropology [5], public health [6], criminology [7], and even accountancy [8]. Life history theory has recently been characterised as offering the unifying meta-theory that the social sciences currently lack [9]. Thus, life history theory is a potentially key inter-disciplinary bridge, connecting diverse human-focussed disciplines to evolutionary theory, and hence grounding human-specific knowledge in a general framework that applies to all living organisms.

As scientific areas grow, they often fission into partially isolated sub-areas. That is, researchers in a field spontaneously form fluid and informal sub-groups, and rely on and credit the ideas generated within their group more than those of surrounding ones. These sub-groups are akin to ‘demes’ in population biology: partially genetically isolated sub-populations that can form the basis for local adaptation [10]. Demic structure in science is not necessarily a bad thing: once the demes are well enough separated, theories and assumptions can evolve rapidly within each deme, generating conceptual diversity and adapting to the specific phenomena that deme deals with [10–12]. However, just as in population biology, the existence of demic structure raises classification problems: when are we dealing with locally-adapted varieties of the same kind of thing, and when are we giving the same name to what have in fact become different kinds of thing? We aimed to understand whether, as life history theory has expanded its range, it has remained a coherent, connected research programme; or whether it has developed deep demic divisions; and if there are such divisions, where they lie.

The emergence of demic structure can be detected in citation patterns, which can be studied quantitatively using bibliometrics. Bibliometrics originated in the early twentieth century [13], with key methodological developments in the 1960s [14–16]. Its more recent development has been facilitated by the availability of comprehensive searchable databases, such as Web of Science (Clarivate Analytics, http://www.webofscience.com). Two important and related goals of bibliometrics are the construction of maps—that is, representations of which parts of a literature are most closely connected to one another [17]; and the detection of clusters—that is, subsets of a literature that are more closely connected internally than they are to parts outside the cluster [18]. Clusters detected in the literature imply demic structure in the research field. Map-making and cluster analysis have allowed researchers to address questions about the structure of science as a whole [19], as well as of individual disciplines [20] or inter-disciplinary research areas [21].

Construction of maps and analysis of clusters require a choice of nodes, and a choice of connections. Nodes can be individual publications, journals, subject categories, or authors. Here, we selected individual publications as the nodes, since using life-history theory is a property of an individual work, not of a journal or an author. To identify connections amongst works, we used bibliographic coupling [22]. Two works are bibliographically coupled if there exists some third work that they both cite; and the more such cited works in common there are, the stronger the coupling. Thus, bibliographic coupling provides a measure of the extent to which different works draw on the same set of intellectual influences. For a set of published works, the matrix of all the possible pairwise bibliographic coupling strengths can be used to make a distance-based map that places works that have more similar lists of cited references closer together, and also for the detection of clusters of works whose cited reference lists are particularly similar to one another. Tools for doing this are freely available using the VOS Viewer software [23].

The origins of life history theory lie in the development of formal mathematical models of the fitness consequences of variation in life history traits [for influential early sources, see 24, 25]. The body of such models was classically reviewed by Stearns [26] and Roff [27,28], and expanded by the development of state-dependent models in the 1980s and 1990s [29,30]. So dominated was the field by formal modelling that in 1976, Stephen Stearns was able to complain that the field had too much theory and not enough data [24]. In 1980, he estimated that the proportion of papers on life history problems based solely on mathematical models was stabilizing at about 30%, which he described as ‘a healthy balance’ between theory and data [31]. We aimed to ascertain whether this balance has changed in more recent times.

From the comments described above, it is clear that what Stearns [24] meant by ‘theory’ was the practice of formal mathematical modelling of fitness in relation to life history traits, rather than any particular empirical claim that might arise from such models. In psychology and most social sciences, the term ‘theory’ often refers to broad descriptive and interpretive frameworks, not necessarily based on mathematical formalizations. Thus, we expect the proportion of life history papers using formal models to be lower in psychology and social science than in biology. This raises the question of what, if not formal modelling, the psychological and social-science work interprets the term ‘life history theory’ to mean. Our impression [from, for example, 1] is that the core idea most often appealed to under the banner of life history theory is that of the ‘fast-slow continuum’ (or ‘fast’ and ‘slow’ strategies): the descriptive generalization that variation between species or between individuals can be organized onto a principal axis from early reproduction, high reproductive effort and short life at one end, to late reproduction, low reproductive effort and long life at the other. Thus, we searched for appeals to the idea of the fast-slow continuum in the bibliometric data, to characterise how these are distributed across research areas and time.

## Methods

We searched Web of Science (Core Collection) on November 12^th^, 2018 for the term ‘life history theory’ in ‘topic’ (for discussion and justification of this search term, see appended Supporting Information, section 1). A ‘topic’ search includes occurrences of the search term in title, abstract, keywords, and keywords plus. There were 1841 records meeting the search criterion. Web of Science searches are not sensitive to hyphenation, and thus our search also returned occurrences of ‘life-history theory’. The Web of Science database goes back to 1970. Records were downloaded as text files for bibliometric coupling analysis in VOS Viewer [23] and as a bibtex database for ancillary analysis using R package ‘bibliometrix’ [32]. The median publication year of papers in the dataset was 2010. Since we were interested in how the structure of the literature has changed over time, we divided the dataset into two parts: publications appearing up to and including 2010 (n = 911), and those appearing later than 2010 (n = 930). We also analysed the whole dataset together (Supporting Information, section 2), and using a finer-grained division into four time periods (Supporting Information, section 3).

An alternative to bibliographic coupling is co-citation analysis. In co-citation analysis, the nodes on the resulting maps are not the papers found by the literature search, but the papers *cited by* the papers found in the literature search (i.e. those papers that, in bibliographic coupling, determine the proximity). Maps using co-citation analysis were generally similar to those presented here (Supporting Information, section 4). However, we preferred bibliographic coupling to co-citation for our main analysis, as a number of the nodes in the co-citation maps were works that were not themselves about life history theory (for example, papers on statistical or measurement methods). Moreover, bibliographic coupling has recently been argued to produce slightly more accurate maps of a research area [33].

We created maps were based on the 500 documents with the greatest link strength. We used the fractional counting method, which equalizes the weight given to each paper [34], including all papers returned by the search regardless of whether they had been cited. We set the cluster resolution parameter at 0.80, with minimum cluster size 20. The choice of cluster resolution is a research judgment, designed to produce an interpretable number of clusters. In the Supporting Information, section 5, we report the effects of varying the cluster resolution in steps from 0 to 1. To normalize the maps for visualization, we used the fractionalization method (for full details of the options available in VOS Viewer, see [23]).

Having created the maps, in order to reveal the content of the identified clusters, we selected a sample of 10 papers that had been assigned to each cluster (see Supporting Information, section 6, for lists of the sampled papers). We chose papers that either presented empirical data or a formal model, and that mentioned life history theory more than just as a keyword. Among papers meeting these criteria, we selected the 10 with the highest total link strength from each cluster. For the 10 sample papers from each cluster, we downloaded the full electronic version and scored four variables: the taxa of organism from whom empirical data is reported; the presence of a formal model; the general topic; and whether they specifically mention the ‘fast-slow continuum’ or ‘fast/slow life history strategies’. Based on the scores for the sample papers on these variables, we assigned names to the clusters. In addition, we tabulated the 10 references most frequently cited by the papers in each cluster. We also counted how many of these top-cited references presented formal models.

For each cluster in each time period, we calculated indices of connection. Two papers are connected in bibliographic coupling analysis if there is at least one reference that both papers cite. For each cluster, we calculated the number of possible links that could exist (*k_i_* × *k_j_* for two different clusters with *k_i_* and *k_j_* papers in them respectively, but *k_i_* × *(k_i_-1)* for links amongst papers in the same cluster). We then computed how many of those possible links actually existed. This gives an index that ranges from 0 (there are no bibliographic links between the two clusters) to 1 (every paper in the first cluster shares a common citation with every paper in the second). As well as the absolute value of the connection index, we also computed two measures of its inequality. First, we calculated the Gini coefficient of each cluster’s connection indices. A connection Gini of 1 for a cluster means that all of its connections are within-cluster and none to other clusters. A connection Gini of 0 means that links from that cluster are perfectly equally distributed across all clusters (0 is impossible in practice because no clusters would be detected without some inequality in connection). Second, we calculated the within/between connection ratio. This is the ratio of the within-cluster connection index to the mean of the connection indices to all the other clusters.

## Data availability

A comprehensive data archive is available at http://doi.org/10.5281/zenodo.2530683. This includes the raw Web of Science records, the lists of papers in each cluster, the VOS Viewer files and the R code required to reproduce the analyses. Readers can also create interactive high-resolution versions of figure 2 by downloading VOS Viewer from http://www.vosviewer.com/ and using the map and network files provided in the data archive.

**Figure 2.**
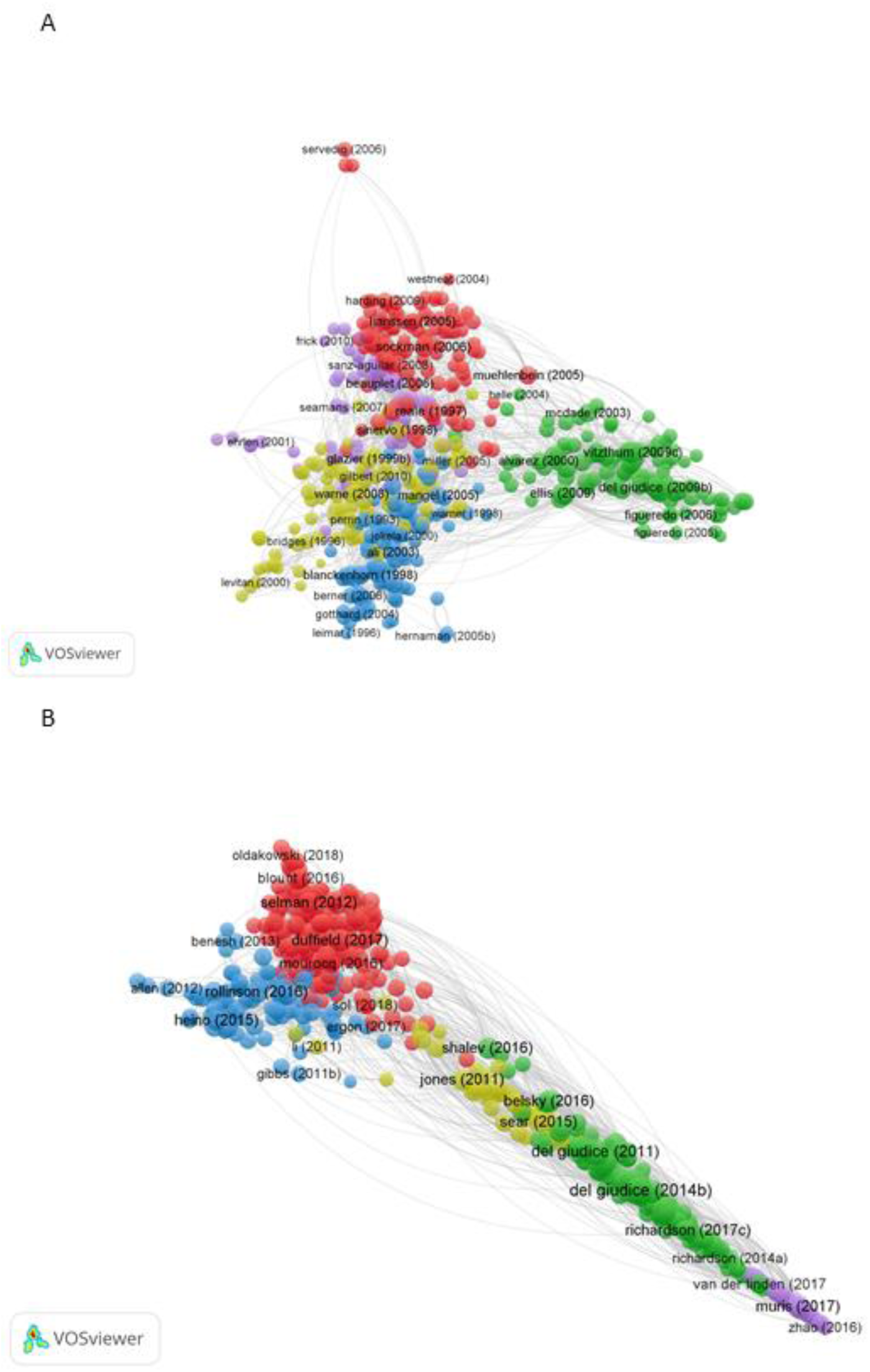
Maps of the life-history theory literature based on bibliographic coupling. A. The literature up to and including 2010. B. The literature published after 2010. For details of parameters used to construct maps, see Methods. Colours are assigned independently to the clusters in the two maps and so shared colour only identifies linkage within but not between time periods.

## Results

The earliest reference to life history theory identified by the search was a paper by Charnov from 1979 [35]. After an initial phase of exponential growth, the number of new papers has grown linearly since around 2005 (figure 1). From 2012 onwards, there were more than 100 new papers every year.

**Figure 1.**
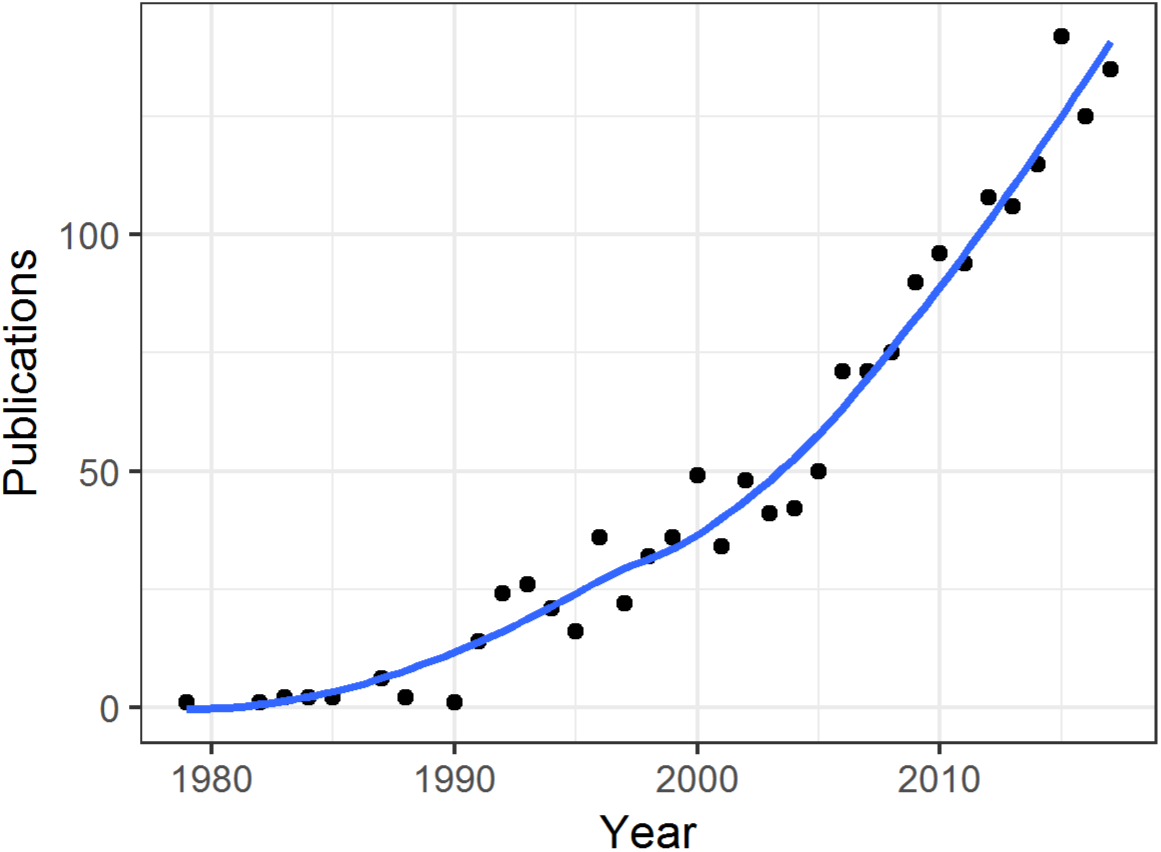
Number of publications found using the topic search term ‘life history theory’ in Web of Science. The year 2018 is not shown in this figure as it was not complete at time of writing. Line represents a loess (locally-estimated non-parametric) fit.

### Structure of the earlier literature (up to and including 2010)

The structure of the earlier literature was radial, with short arms protruding from a central core (figure 2A). There were five clusters. The primary differences between the clusters appeared to be taxonomic (table 1). In our sample papers, cluster A1 featured birds exclusively; A4 featured reptiles in 7/10 sampled papers; and A5 featured non-human mammals in 9/10 sampled papers. Cluster A3 was a mixture of work on invertebrates, fish, and amphibians. Cluster A2 captured all of the research on humans. Some non-human primate data also appeared in the sample papers from cluster A2, often in comparison to humans. Unlike the taxa, the topics of study did not obviously differ strongly by cluster, with fundamental life history questions concerning reproductive effort and its costs, parental investment, and growth, appearing in all clusters. Formal models were found in 16 of the 50 papers sampled in this period, distributed across all clusters except A5. The lists of 10 top-cited references for the five clusters of this time period showed some overlap, with Stearns’ book [26] appearing in the top 10 for every cluster. For all clusters, three or more of the top 10 cited references were sources that presented formal evolutionary models. In this earlier time period, the idea of the ‘fast-slow continuum’ or ‘fast/slow life history strategies’ occurred in four of the 50 sample papers (one in cluster A2, three in A5).

**Table 1.** Data on the clusters identified by the bibliographic coupling analysis, period up to and including 2010.

| Cluster | n | Name | Topics | Taxa | Formal model in sample papers belonging to cluster | Fast-slow axis | References most cited by papers belonging to cluster | Formal model in ten sources most cited by papers belonging to cluster |
| --- | --- | --- | --- | --- | --- | --- | --- | --- |
| A1 | 120 | Birds | Reproductive effort, ageing | Birds 8/8 | 5 / 10 | 0 / 10 | 1. *ELH-S*; 2. *ELH-R*; 3. Drent, R. H. (1980) *Ardea* 68: 225; 4. Clutton-Brock, T. C. (1991). *Evolution of Parental Care* (Princeton); 5. Deerenberg, C. (1997). *Proc. Roy. Soc. B* 264: 1021; 6. Linden, M. (1989). *Trends. Ecol. Evol.* 4: 367; 7. Martin, T.E. (1987). *Ann. Rev. Ecol. Systematics* 18: 453; 8. Lack, D. (1968). *Ecological Adaptations for Breeding in Birds* (Chapman Hall); 9. Lessells, C. M. (1991) in: *Behavioural Ecology: An Evolutionary Approach* (Blackwell); 10. Sheldon, B. C. (1996). *Trends. Ecol. Evol.* 11: 317 | 3 / 10 |
| A2 | 115 | Humans | Reproductive effort, ageing, childhood psychosocial experience | Humans 10 / 10<br>Non-human primates 1/10 | 3 / 10 | 1/10 | 1. *ELH-S*; 2. Belsky, J. (1991). *Child Dev.* 62: 647; 3. Charnov, E. L. (1993). *Life History Invariants* (Oxford); 4. Chisholm, J.S. (1993). *Curr. Anthro.* 34: 1; 5. Kaplan, H.S. (2000). *Evol. Anthro.* 9: 156; 6. Hill, K. (1996). *Ache Life History* (Routledge); 7. Hawkes, K. (1998). *PNAS* 95: 1336; 8. Hill, K. (1999). *Ann. Rev. Anthro.* 28: 397; 9. *ELH-R*; 10. Bogin, B. (1999). *Patterns of Human Growth* (Cambridge). | 4 / 10 |
| A3 | 109 | Invertebrates and others | Growth, maturation, theory | Invertebrates 3 / 6<br>Fish 2 / 6<br>Amphibians 1 / 6 | 6 / 10 | 0 / 10 | 1. *ELH-R*; 2. *ELH-S*; 3. Abrams, P.A. (1996). *Am. Nat.* 147: 381; 4. Charlesworth, B. (1980). *Evolution in Age-Structured Populations* (Cambridge); 5. Arendt, J.D. (1997). *Q. Rev. Biol.* 72: 149; 6. Cole, L.C. (1954). *Q. Rev. Biol.* 29: 103; 7. Stearns, S.C. (1992). *Evolution* 40: 893; 8. Fisher, R.A. (1930). *The Genetical Theory of Natural Selection* (Oxford); 9. Stearns, S.C. (1976). *Q. Rev. Biol.* 51: 3; 10. Atkinson, D. (1994). *Adv. Ecol. Res.* 25: 1. | 8 / 10 |
| A4 | 93 | Reptiles and others | Reproductive effort, parental investment | Reptiles 7 /10<br>Fish 2 / 10<br>Invertebrates 2 / 10<br>Non-primate mammals 1 / 10<br>Non-human primates 1 / 10 | 2 / 10 | 0 / 10 | 1. *ELH-R*; 2. *ELH-S*; 3. Reznick, D. (1985). *Oikos* 44: 257; 4. Smith, C.C. (1974). *Am. Nat.* 108: 499; 5. Gadgil, M. (1970). *Am. Nat.* 104: 1; 6. Williams, G.C. (1966). *Am. Nat.* 100: 687; 6. Houle, D. (1992). *Genetics* 130: 195; 8. McGinley, M.A. (1987). *Am. Nat.* 130: 370; 9. Parker, G.A. (1986). *Am. Nat.* 128: 573; 10. Bell, G. (1986). *Ox. Surv. Evol. Biol.* 3: 83. | 9 / 10 |
| A5 | 63 | Mammals | Reproductive effort, ageing, maturation, fast-slow continuum | Non-primate mammals 9 / 10<br>Non-human primates 2 / 10<br>Birds 1 / 10 | 0 / 10 | 3 / 10 | 1. Burnham, K. P. (1998). *Model Selection and Multimodel Inference* (Springer); 2. LeBreton, J.D. (1992). *Ecol. Monographs* 62: 67; 3. *ELH-S*; 4. Brownie, C. (1993). *Biometrics* 49: 1173; 5. Caswell, H. (2001). *Matrix Population Models* (Oxford); 6. Cole, L.C. (1954). *Q. Rev. Biol.* 29: 103; 7. Clutton-Brock, T. C. (1982). *Red Deer* (Chicago); 8. Gaillard, J.M. *Ann. Rev. Ecol. Systematics* 31: 367; 9. Cam, E. (1998) *Ecology* 79: 2917; 10. Cam, E. (2002). *Am. Nat.* 159: 96. | 3 / 10 |

Notes: Top cited references are in descending order of citations. First author name only is shown. *ELH-S*: Stearns, S.C. (1992). *The Evolution of Life Histories* (Oxford); *ELH-R*: Roff, D.H. (1992/2002). *The Evolution of Life Histories* (Chapman and Hall). Note that taxa numbers do not necessarily add to 10 as some papers contain only models, and others contain data from multiple taxa.

### Structure of the recent literature (post 2010)

The recent literature had a more linear structure, with relatively few direct bibliographic links between one end and the other (figure 2B). Again, there were five clusters (table 2). These separated initially on taxonomic lines: birds and non-primate mammals (B1); fish, plants, and invertebrates (B3); and humans (B2, B4, and B5). Within the human clusters, B2 represented developmental and personality psychology, focussing on individual variation and the impacts of childhood psychosocial experience. Cluster B4 represented the research of biological and evolutionary anthropologists, often including data from small-scale societies, and in some cases non-human primates. Finally, a small cluster, B5, represented a more specific subset of human personality psychology, concerned in particular with the ‘dark triad’ traits (Machiavellianism, narcissism, and psychopathy). The three human-focussed clusters consisted of a total of 225 papers (45% of the total). This compares to 115 (23%) of papers belonging to the sole human cluster A2 in the earlier time period.

**Table 2.** Data on the clusters identified by the bibliographic coupling analysis, post 2010.

| Cluster | N | Name | Topics | Taxa | Formal model in sample papers belonging to cluster | Fast-slow axis | References most cited by papers belonging to cluster | Formal model in ten sources most cited by papers belonging to cluster |
| --- | --- | --- | --- | --- | --- | --- | --- | --- |
| B1 | 184 | Birds and mammals | Reproductive effort, ageing, parental investment | Birds 7/10<br>Non-primate mammals 4/10 | 0 / 10 | 2/10 | 1. *ELH-S*; 2. Burnham, K. P. (1998). *Model Selection and Multimodel Inference* (Springer); 3. Harshman, L.G. (2007). *Trends. Ecol. Evol.* 22: 80; 4. Alonso-Alvarez, C. (2004). *Ecol. Letters* 7: 363; 5. Clutton-Brock, T.H. (1984). *Am. Nat.* 123:212; 6. Bergeron, P. (2011). *Funct. Ecol.* 25: 1063; 7. Clutton-Brock, T. C. (1991). *Evolution of Parental Care* (Princeton); 8. Garratt, M. (2011). *Proc. Roy. Soc. B* 278: 1098; 9. Van Noordwijk, A.J. (1986). *Am. Nat.* 128: 137; 10. Williams, G.C. (1966). *Am. Nat.* 100: 687. | 3 / 10 |
| B2 | 155 | Human developmental/ personality | Childhood psychosocial experience in relation to maturation and psychological traits | Humans 10/10 | 0/10 | 10/10 | 1. Ellis, B.J. (2009). *Hum. Nat.* 20: 204; 2. Belsky, J. (1991). *Child Dev.* 62: 647; 3. Figueredo, A.J. (2006). *Dev. Rev.* 26: 243; 4. Chisholm, J.S. (1993). *Curr. Anthro.* 34: 1; 5. Ellis, B.J. (2004). *Psych. Bull.* 130: 920; 6. Belsky, J. (2012). *Dev. Psych.* 48: 662; 7. Figueredo, A.J. (2004). *Soc. Biol.* 51: 121; 8. Del Giudice, M. (2009). *Behav. Brain. Sci* 32: 1; 9. Brumbach, B.H. (2009). *Hum. Nat.* 20: 25; 10. Figueredo, A.J. (2005). *Pers. Ind. Diff.* 39: 1349. | 0 / 10 |
| B3 | 90 | Fish and others | Reproductive effort, growth, ageing, parental investment, energetics | Fish 4/7<br>Plants 2/7<br>Invertebrates 1/7 | 3/10 | 0/10 | 1. *ELH-S*; 2. *ELH-R*; 3. Bell, G. (1980). *Am. Nat.* 116: 45; 4. Bernado, J. (1996). *Am. Zool.* 36: 216; 5. Smith, C. C. (1974). *Am. Nat.* 108: 499; 6. Allen, R.M. (2008). *Am. Nat.* 171: 225; 7. Berkeley, S.A. (2004). *Ecology* 85: 1258; 8. Caswell, H. (2001). *Matrix Population Models* (Oxford); 9. Conover, D. H. (2002). *Science* 297: 94; 10. Heino, M. (2002). *Evolution* 56: 669. | 5 / 10 |
| B4 | 40 | Human evolutionary | Reproductive effort, growth, mortality, immunity, alloparenting, skill acquisition | Humans 8/10<br>Non-human primates 3/10 | 0/10 | 2/10 | 1. Kaplan, H.S. (2000). *Evol. Anthro.* 9: 156; 2. Hawkes, K. (1998). *PNAS* 95: 1336; 3. Hill, K. (1996). *Ache Life History* (Routledge); 4. *ELH-S*; 5. Bogin, B. (1997). *Yearb. Phys. Anthro.* 40: 63; 6. Charnov, E.L. (1993). *Evol. Anthro.* 1: 191; 7. Gurven, M. (2006). *Proc. Roy. Soc. B* 273: 835; 8. Hamilton, W.D. (1966). *J. Theor. Bio.* 12: 12; 9. Kaplan, H.S. (2002). *PNAS* 99: 10221; 10. Kuzawa, C.W. (2012). *Curr. Anthro.* 53: 369. | 4 / 10 |
| B5 | 29 | Dark triad personality | 'Dark triad' personality traits (Machiavellianism, narcissism, and psychopathy) | Humans 10/10 | 0/10 | 10/10 | 1. Jonason, P.K. (2009). *Eur. J. Pers.* 23: 5; 2. Jonason, P.K. (2010). *Hum. Nat.* 21: 428; 3. Paulhus, D.L. (2002). *J. Res. Pers.* 36: 556; 4. Jonason, P.K. (2010). *Psychol. Assessment* 22: 420; 5. Christie, R. (1970). *Studies in Machiavellianism* (Academic Press); 6. Figueredo, A.J. (2006). *Dev. Rev.* 26: 243; 7. Jonason, P.K. (2012). *Rev. Gen. Psych.* 16: 192; 8. Jonason, P.K. (2010). *Pers. Ind. Diff.* 49: 611; 9. Jonason, P.K. (2012). *Pers. Ind. Diff.* 52: 521; 10. Jones, D.N. (2011). *Pers. Ind. Diff.* 51: 679. | 0 / 10 |

Notes: Top cited references are in descending order of citations. First author name only is shown. *ELH-S*: Stearns, S.C. (1992). *The Evolution of Life Histories* (Oxford); *ELH-R*: Roff, D.H. (1992/2002). *The Evolution of Life Histories* (Chapman and Hall). Note that taxa numbers do not necessarily add to 10 as some papers contain only models, and others contain data from multiple taxa. Note that taxa numbers do not necessarily add to 10 as some papers contain only models, and others contain data from multiple taxa.

The topics studied did not obviously separate clusters B1, B3 and B4; but B2 with its focus on psychological variation and childhood psychosocial experience, and B5 with its more specific focus on the dark triad personality traits, were distinct from the others in topic as well as taxon. Formal evolutionary models were found in just 3 of the 50 papers sampled from the post-2010 literature (all in cluster B3). Unlike the earlier time period, there was no item common to 10 most cited reference list of all the clusters. The classic formal models [26] continued to be prominent in the top-cited references of clusters B1, B3 and B4. The top-cited lists from clusters B2 and B5 contained no references to non-human work or to any formal evolutionary models. The ideas of the ‘fast-slow continuum’ or ‘fast and slow strategies’ were much more frequent in the later than the earlier time period, being found in 24 of the 50 sampled papers. This was due to the universal (10/10) allusion to these ideas in the sample papers from clusters B2 and B5. However, mention of the ideas remained rare outside of these two clusters.

### Connections within and between clusters

The connection index in the earlier time period was 0.56 overall (i.e. 56% of possible bibliographic links actually existed). Links were well distributed across clusters (figure 3A): indeed, the papers from all clusters were only modestly better linked to other papers within their cluster than to papers outside their cluster. Reflecting this, the Gini coefficients for the inequality of connections across clusters were low (A1, 0.08; A2, 0.10; A3, 0.09; A4, 0.10; A5, 0.09), and the within/between connection ratios were only slightly greater than 1 (A1, 1.23; A2, 1.48; A3, 1.30; A4, 1.32; A5, 1.34). In the later time period, the connection index overall was 0.35. The distribution of links across clusters was much more uneven (figure 3B). Correspondingly, the Gini coefficients of connection were higher than the earlier time period, especially for cluster B5 (B1, 0.39; B2, 0.29; B3, 0.39; B4, 0.36; B5 0.80), as were the within/between link ratios (B1, 2.16; B2, 2.25; B3, 2.07; B4, 2.34; B5 7.27). Cluster B5 was only connected to the rest of literature via B2; its connection indices to all other clusters were close to zero. Cluster B2 had rather low connection indices (less than 0.30) to the two non-human clusters B1 and B3. This means that there is relatively little overlap between the citation lists of the papers in cluster B2 and B5 and the citation lists of papers on non-human organisms.

**Figure 3.**
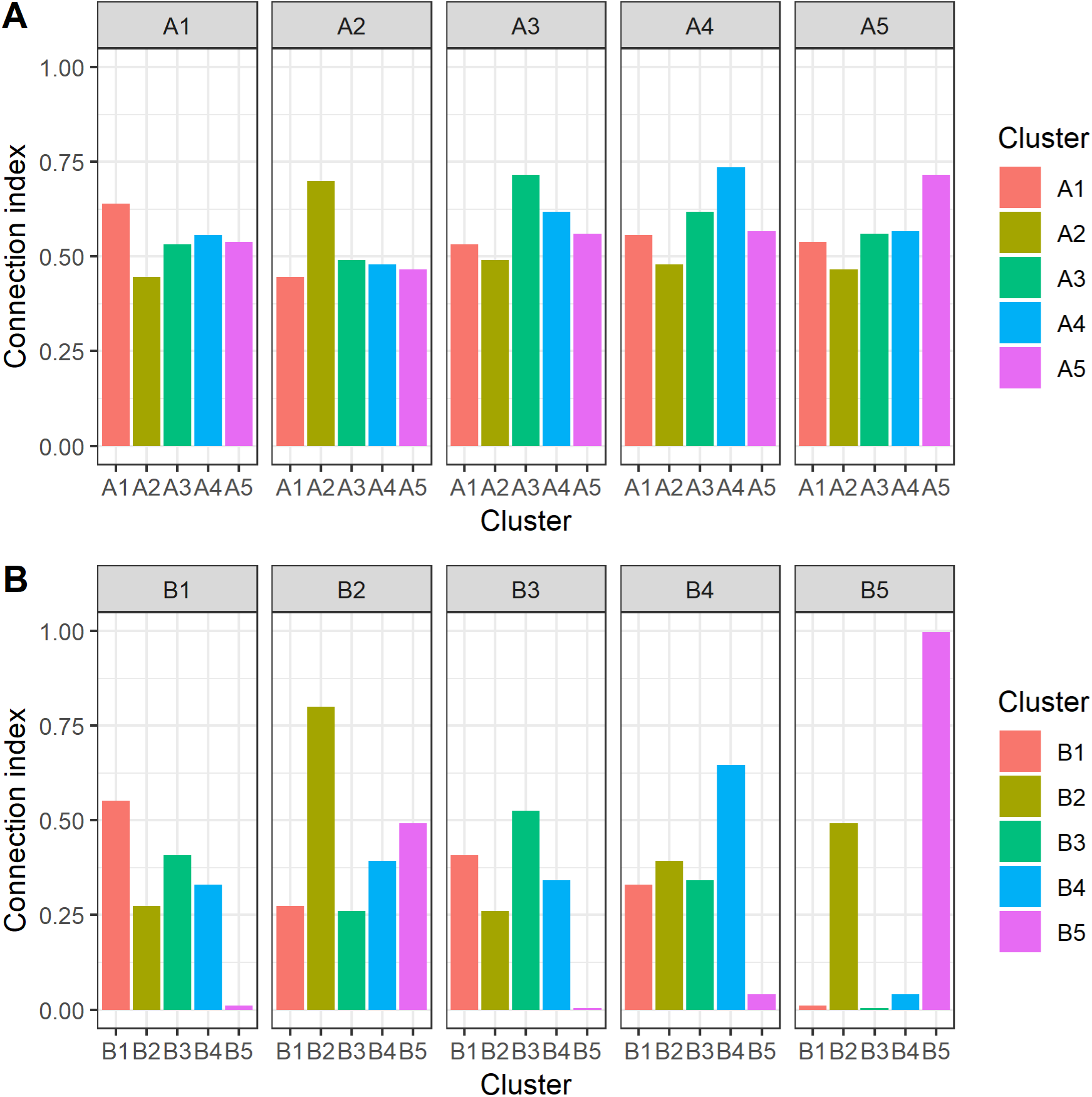
Indices of bibliographic connection between papers in the different clusters in the period up to and including 2010 (A) and the period since 2010 (B). An index of 1 means that all papers are connected by a shared cited reference, and 0 means there are no shared citations between any of the papers in the two clusters.

## Discussion

Using an approach based on bibliographic coupling, we documented the changing structure of the life history theory literature over time. Our cluster analysis suggests that the field has always been divided to some extent on taxonomic lines, with work on birds, mammals, other organisms, and humans tending to show some separation in the sources they cite. Between the early period (up to and including 2010) and the more recent literature, the proportion of papers on humans greatly increased, and the human literature has fractionated into distinct literatures from evolutionary anthropology and from different areas of psychology. (In fact, the basic differences between the various human approaches existed from the earliest time period, but they were not differentiated enough to be detected as separate clusters until more recently; see Supporting Information, section 3). The three human clusters—evolutionary anthropology, developmental/personality psychology, and dark triad—lie at progressively greater bibliometric distance from life history theory as it is practiced in ecology and evolutionary biology. The internal structure of the human literature is approximately but not exactly captured by Black et al.’s [36] distinction between ‘bio-demographic’ studies (i.e. focused on growth and reproduction) and ‘psychological’ studies (i.e focussed on behavioural or cognitive traits such as prosociality, personality and religiosity, that are argued to be related to life history strategy): though all ‘psychological’ approaches are in clusters B2 and B5, some ‘bio-demographic’ outcomes such as age at menarche are also studied in cluster B2 [e.g. 37].

Though we detected five clusters in both time periods, there were several lines of evidence that the literature has become more fragmented. Up to 2010, the probability of two papers sharing a common cited reference was only modestly higher (around 30%) if they belonged to the same cluster than if they belonged to different ones. Reflecting this, the Gini coefficients of the inter-cluster connection index were low, indicating that every part of the literature was nearly equally connected to every other part through shared citations, and there was overlap in the most commonly cited references of all clusters. In the post-2010 period, the overall probability of any two papers sharing a cited reference was lower, and links were much more concentrated within clusters (the probability of a shared citation being at least 100% greater for two papers of the same rather than different clusters). The lower overall probability of sharing a common citation was expected: as the literature has become so much larger, and citation lists are finite, the expected amount of overlap should decrease. However, the increasing dominance of within-cluster citation over between-cluster citation is not an inevitable concomitant of the growth in the size of the literature. Instead, this suggests formation of separate ‘demes’ of research all referring to their content as life history theory. This conclusion is backed up an ancillary analysis of the effects of gradually increasing the sensitivity of the cluster detection algorithm (Supporting Information, section 5). In the earlier time period, the resolution can be up to 0.26 before any discrete clusters are detected, whereas in the more recent time period, a cluster division is detected with a resolution of 0.15. The initial division in both cases is between human and non-human research.

The shape of the map of the literature up to and including 2010 was basically radial. This means that every cluster has a zone of proximity to every other cluster, at the centre of the map. To document this phenomenon another way, the list of 10 top-cited references in every A cluster contained common theoretical resources, notably the books by Stearns [26] and Roff [27,28]. By contrast, the post-2010 map was rather linear. This means that the papers in the clusters at one end are only connected to those at the other via a series of intermediate links: dark triad research is linked to research in developmental and personality psychology, which is linked to research on human evolution, which is linked to non-human research. Reflecting this, there were no sources found in the 10 top-cited references of clusters B2 and B5 that were found in the top-cited list of any other cluster.

The existence of discrete clusters is not necessarily a bad thing for a research programme. It is quite understandable that researchers are particularly likely to cite prior results (or methodologies) using their study species or a closely related one. This facilitates local adaptation of methods and ideas to a particular application. Some degree of clustering may be advantageous in terms of the generation of conceptual diversity [10–12]. Moreover, in some of the post-2010 work, particularly cluster B5, life history theory is not the central theme, but an ancillary resource alluded to in framing the question, sometimes mentioned only in keywords. However, very deep clustering does pose questions of nomenclature: how helpful is it that the different clusters all use the same term, life history theory? If the probability of shared citation across clusters becomes sufficiently low, conceptual evolution becomes independent: at this point, the term ‘life history theory’ may no longer have a unique referent. Claims about the successes, flaws, predictions or findings of life history theory then become problematic: one set of ideas can end up receiving credit or responsibility for the outputs of a different one.

The striking division within our post-2010 sample papers is in allusion to the concept of a ‘fast-slow’ continuum, or ‘fast’ and ‘slow’ life history strategies. Appeal to this concept is universal in papers from clusters B2 and B5. Indeed, within these clusters, the fast-slow continuum is presented as the central defining idea that individuates life-history theory as opposed to other approaches. In fact, though, the fast-slow continuum construct is rarely mentioned elsewhere in the literature (4 of 30 of the sampled papers from the other B clusters; [38], a paper on birds, is a rarity in framing life history theory in similar terms to the papers from clusters B2 and B5). Moreover, the fast-slow continuum concept, at least under that name, was rare in the data up to and including 2010. The ‘fast-slow’ terminology originates in empirical research on cross-species patterns of covariation in multiple life history traits [39,40]. As Oli [40] specifies, the fast-slow concept thus arose not from life history *theory* (in the sense of formal modelling), but as an inductive generalization from comparative *data*. Clusters B2 and B5 extend the idea of a fast-slow continuum underlying cross-species variation to within-species variation; extend the idea from life history traits to behavioural and personality variables; and (in B2 especially) are concerned with developmental plasticity in life history strategies.

Biologists working on non-human animals have pursued ideas similar to those found in clusters B2 and B5. They tend to do so, however, under the label ‘pace of life’ or ‘pace of life syndrome’ [for recent and contrasting reviews of this area, see 41,42]. Pace of life research is typically presented as distinct from life history theory: a Web of Science search on ‘pace of life’ produces over 400 records, 95% of which are not found by searching for ‘life history theory’ (see Supporting Information section 3). This separation may be because there is as yet little formal modelling of the evolutionary basis of the pace of life idea [43]. This means pace of life could become more strongly linked to life history theory in the future if the relevant evolutionary models are developed. The central elements of the pace of life paradigm are: that a fast-slow continuum might exist between individuals as well as between species or populations; that this single continuum might organize variation in suites of different traits, including behavioural, personality, and physiological traits; and that the determinants and consequences of being at different positions along such a continuum should be studied [44]. Réale et al. [44] point out that the original source of the pace of life idea is the 1970s idea of r-versus K-strategists. Black et al. [36] make exactly the same point with respect to the human fast-slow psychological paradigm; indeed, early names for that paradigm were ‘r-K’ or ‘differential K’ theory [45,46]. Thus, we would argue that the human research in clusters B2 and B5 of our dataset belongs more logically with the non-human pace of life research than it does with life history theory as that term is typically used in non-human biology. However, at present, clusters B2 and B5 and the pace of life paradigm are not making much reference to one another. The three most cited pace of life papers [44,47,48] are cited, respectively, 5, 0 and 3 times by the papers of cluster B2, and 0, 0 and 0 times by the papers of cluster B5. Conversely, we have found only a single paper framing the study of variation in humans explicitly within the pace of life terminology [49].

Our sampling of papers from each cluster suggested there may have been a recent decline in the proportion of the literature that consists of formal models. Up to and including 2010, 16 of 50 papers we sampled presented formal models, a proportion very close to the 30% that Stearns estimated back in 1980 [31]. In the post-2010 sample, we found just three of 50 papers (6%) presenting formal models. This decline was not just due to the absence of formal modelling in the human-focussed clusters: three of 20 papers from the non-human clusters suggests a reduction in the amount of formal modelling there too. It is not clear what might explain this decline.

Researchers may believe that the fundamental theory has already been thoroughly mapped out, and what remains is to test its assumptions and predictions, or elucidate the proximate mechanisms underlying trade-offs. Alternatively, the growth in annual productivity of the research programme may be due to more empiricists being attracted to draw on the theoretical ideas, with the absolute number of theoreticians remaining constant, producing a proportionate decline in theory papers. These remain speculations on our part.

It is possible that the decline in formal modelling, or in direct reliance on the results of such modelling, has exacerbated fragmentation. Verbal statements can readily mutate through qualitative interpretation and memorial bias, so that their transmission resembles a game of telephone. For example, citations in papers are quite often (perhaps 15% of instances) used in support of statements that are substantively different from, absent from, or even contradictory to, the statements in the paper cited [50,51]. When theory is formalized in mathematical models, however, it may be transmitted with greater fidelity. Thus, formalization may play a key role in stabilizing the core of a body of theory. Consider for example Newtonian mechanics, or the modern synthesis version of evolutionary theory: whatever their need for amendment due to the discovery of new phenomena, scientists at least all agree what these bodies of theory consist of. We would predict that research programmes based closely on formal theory will tend to remain more stable and conceptually unified, and that a shift to more verbally-based theory will also lead to more rapid conceptual change and greater fragmentation. These predictions could be tested in a future study.

In conclusion, we have documented the evolution over time of the literature appealing to the concept of life history theory. This literature has grown rapidly and increasingly incorporated more psychological and social-science research. As it has grown, its internal differentiation has become more marked. It may be incipiently speciating into a part concerned with the fast-slow continuum in humans, a part we argue is conceptually allied with the pace of life hypothesis, and a body of work that shares the original concerns of life history theory, drawing more heavily on formal modelling.

## Acknowledgements

This project has received funding from the European Research Council (ERC) under the European Union’s Horizon 2020 research and innovation programme (grant agreement No AdG 666669, COMSTAR, to DN); and by grants from the Netherlands Organization for Scientific Research (016.155.195), the James S. McDonnell Foundation (220020502), the Jacobs Foundation (2017 1261 02), and the Robert Wood Johnson Foundation (73657) to WEF.

## Supporting Information

### Section 1. ‘Life history theory’ and alternative search terms

We based our study on the topic search term ‘life history theory’ because we were interested in the role of theory in particular (rather than empirical effects or techniques) in bridging between different disciplines. Moreover, the phrase ‘life history theory’ is very often used in framing papers, with the implication that it identifies a particular unified research programme. However, it is clear that this search term does not capture all of the literature commonly thought of as contributing to that particular research programme. There are some classic early works now thought of as canonical life history theory [1,2] that are not captured by the search. Stearns [3] estimated that there had been 52 papers on life history evolution by 1980, whereas our search captured only one [4]. Our strategy therefore under-samples the early theoretical work. Our sample is therefore best thought of as the set of literature that employs the exact term ‘life history theory’, rather than the set of literature that contributed to our understanding of the evolution of life history traits.

We also explored other search terms. Using just ‘life history’ returned over 60,000 records, many of which were clearly not related to the research area we wish to study. ‘Life history strategies’ is also widely used, and may in some cases represent an alternate designation for the life history theory research programme. A search for ‘life history strategies’ returned 2753 records, of which 132 were also in the set of records found by ‘life history theory’. Because of our focus on theory, we preferred the term ‘life history theory’ for our main analysis. ‘Life history strategies’ returns a broader, less unified set of records, containing much more research on plants. The map of the ‘life history strategies’ literature has a less easily interpretable structure. For example, figure S1A shows the post-2010 map of ‘life history strategies’, using the same parameter values as figure 2 of the main paper. We manually excluded two papers that are evidently very bibliometrically distant from all others: one on the helminthic infections of eider ducks [5], and another on plankton [6]. The resulting map after these exclusions is shown in figure S1B. In this map, all the human psychology work is concentrated in the bottom right-hand cluster, which has a somewhat linear structure and a separation from the rest of the map. Thus, the literature on ‘life history strategies’ rather than ‘life history theory’ still confirms the relative separation of the human psychological research from comparable research on other taxa.

**Figure S1.**
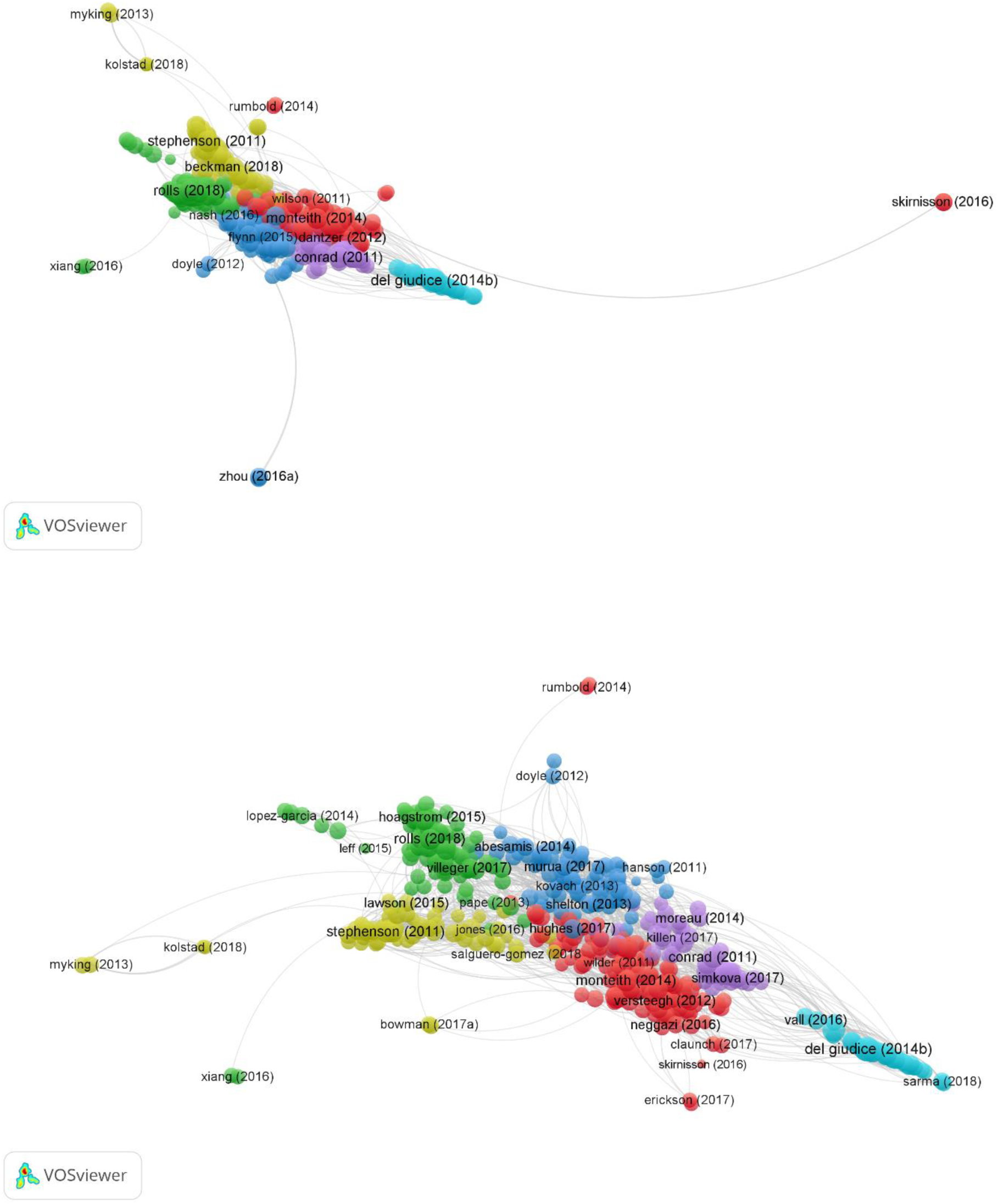
Bibliographic coupling maps of the literature on ‘life history strategies’ post-2010, using the same parameters as the maps in figure 2 of the main paper. A. Original map. B. With manual exclusion of extreme outliers. The bottom right cluster consists of research on humans. We also searched Web of Science for ‘pace of life’ (see Discussion of main paper). This returned 407 hits, all only 22 were also in the ‘life history theory’ set. The papers in the ‘pace of life’ set tend to draw on the ‘fast-slow continuum’ and related ideas. For example, 123 of the 407 mention ‘fast’ in their title or abstract, and 113 mention ‘slow’. As we argue in the main paper Discussion, the research programme of clusters B2 and B5 more logically belongs with the non-human ‘pace of life’ literature than it does with the literature on life history theory.

### Section 2. Mapping the whole of the literature

For our main analysis, we divided the literature into two time periods, up to and including 2010, and post-2010. If instead we use the whole of the literature for mapping, we obtain a map with a similar general shape to the post-2010 map in the main paper (figure S2). As in the main paper, the human research is concentrated along one end of the line, with the ‘dark triad’ research most distant from the rest of the literature. This analysis produces four clusters using the same cluster resolution as in the main paper. These clusters appear to be: birds and mammals (red on figure S2); fish, insects and other taxa (blue); human evolution (yellow); and human developmental/personality psychology (green). The human developmental/personality psychology cluster amalgamates clusters B2 and B5 of the post-2010 map from the main paper: spatially, these two research areas still separate, but the cluster analysis does not consider them separate clusters in this overall analysis.

**Figure S2.**
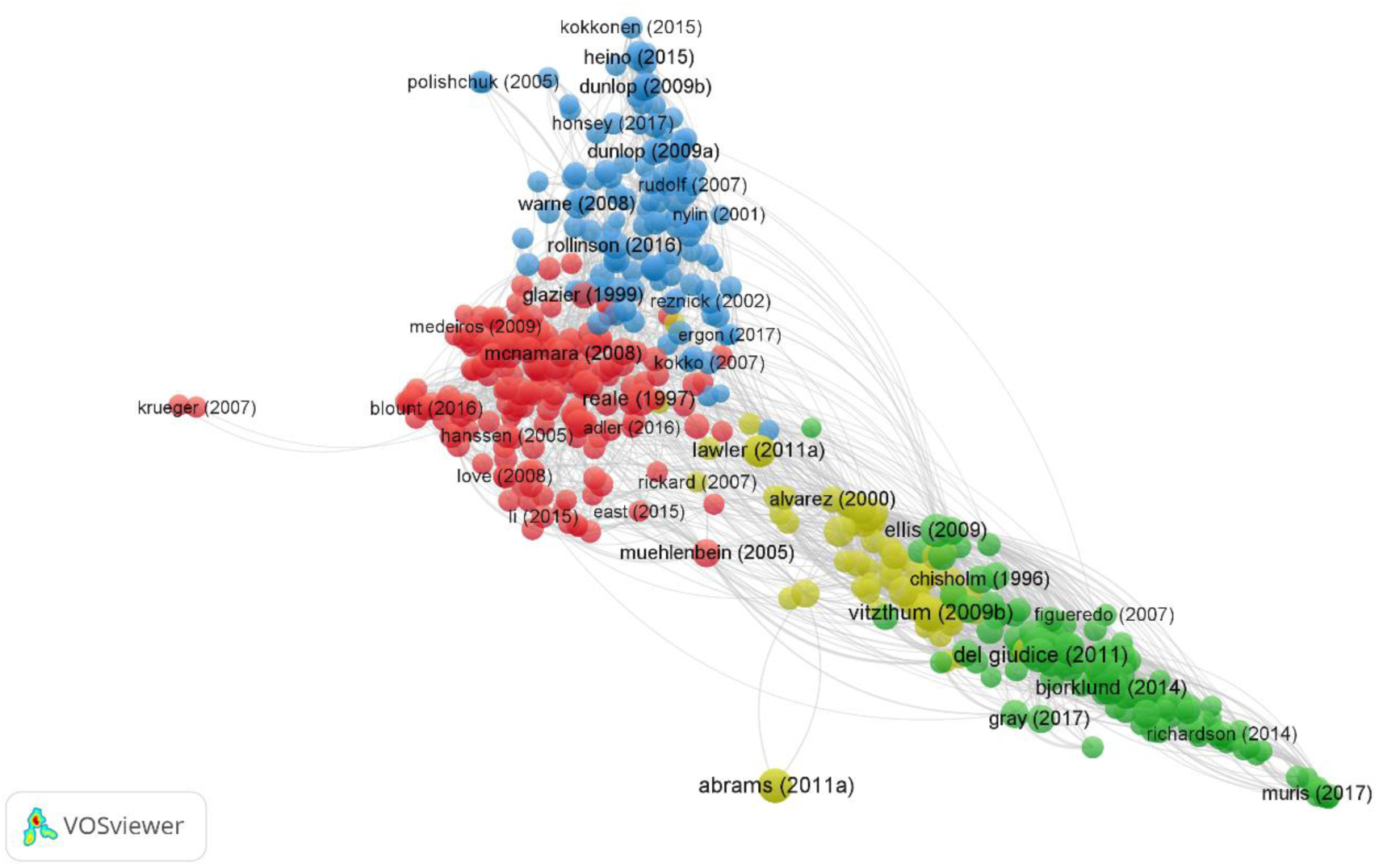
Bibliographic coupling map including both the earlier (up to and including 2010) and recent (post-2010) literature together. Parameter settings are as for figure 2 of the main paper.

### Section 3. Finer-grained division of time

As well as the binary division of time used in the same paper, we also divided the data into four time categories containing roughly equal numbers of records: up to and including 2004 (n = 458); 2005-2010 (n = 453); 2011-2014 (n = 423); and 2015-2018 (n = 508). Figure S3 shows bibliographic coupling maps of each of these four time bins using the same parameter values as the maps in the main paper.

**Figure S3.**
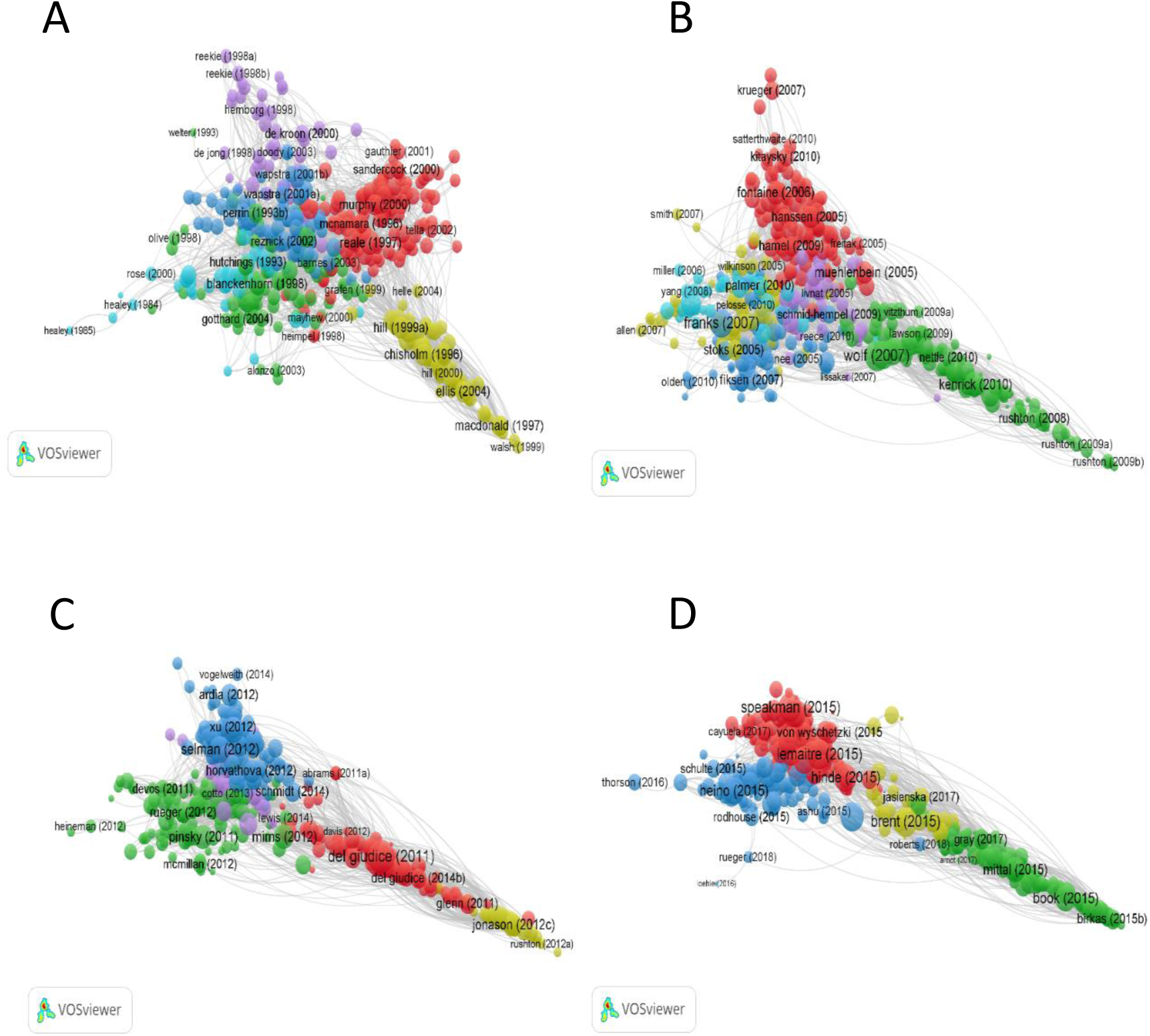
Bibliographic coupling maps for the time periods up to and including 2004 (A); 2005-2010 (B); 2011-2014 (C); 2015-2018 (D). Parameter values are as for figure 2 of the main paper.

The sequence of maps in figure S3 clearly shows the increasingly linearity of the literature over time. It also shows the increasing variegation of the human research: in the first and second time periods, there is only one cluster of human research. In the third time period, the ‘dark triad’ cluster separates from the human development cluster, although the two are combined again in the fourth time period. The division between the human evolution and human developmental/personality psychology is only detected by the clustering algorithm in the most recent time period. However, although not yet marked enough to be detected by the clustering algorithm, the difference between the three kinds of human life history theory research on humans existed from the very earliest time period. For example, in the period prior to 1995, Helle et al. [7], Chisholm [8] and Rushton [9] respectively already exemplified what would later become the three human clusters. The relative positions of these three types of research on the map, and their relative distance from non-human research, are completely consistent across the time periods.

### Section 4. Co-citation maps

An alternative to the bibliographic coupling method we use in the main paper is co-citation analysis. Here, the nodes on the map are the papers cited by the papers found in the literature search. The links are formed by being cited by the same source. Co-citation analysis should produce similar representations of the structure of the field as bibliographic coupling [10]. We repeated our mapping exercise using co-citation analysis instead. We used a minimum number of citations of eight for a node to be included. We also set the cluster resolution parameter to 1.0 instead of 0.8 as for the bibliographic coupling: the number of clusters found at any given resolution was fewer in the co-citation analysis. Other parameter values were as for the bibliographic coupling. The resulting maps were as shown in figure S4.

The same basic generalizations hold as for the bibliographic coupling maps, especially the increasing linearity of structure of the map. In the earlier time period, the clustering identified four rather than five clusters; as in the main paper, all the human research was in one of these. In the later time period, the clustering identified six rather than five clusters. Three of these (the three human clusters, B2, B4 and B5) were identical to the bibliographic coupling clusters. The non-human literature was divided into three clusters here rather than the two in the bibliographic coupling map.

**Figure S4.**
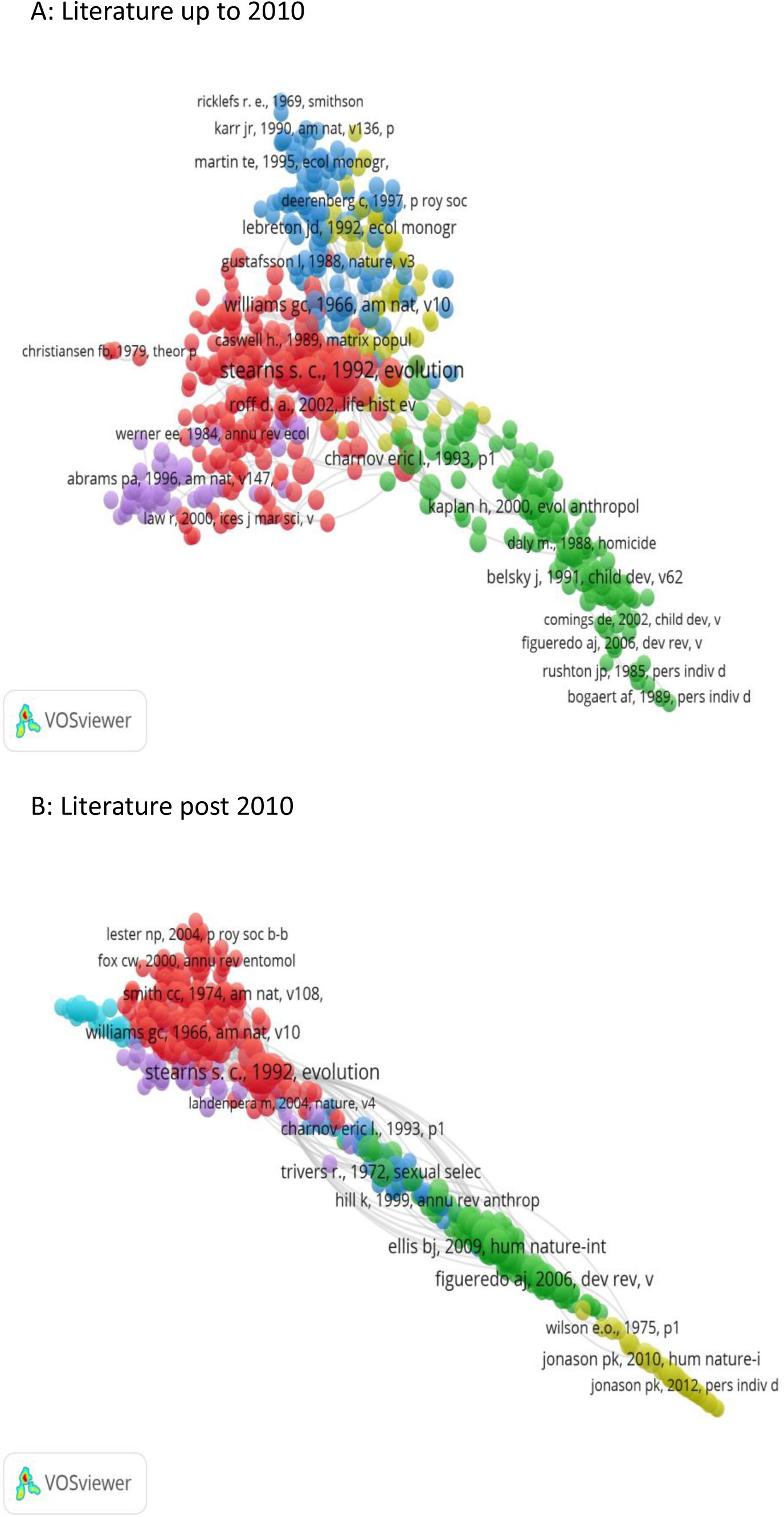
Co-citation maps for the period up to and including 2010 (A) and post-2010 (B). For parameter values see text.

### Section 5. Varying the cluster resolution parameter

The cluster detection is controlled by a resolution parameter (with higher resolution representing greater sensitivity to clustering). We chose 0.80 for the main analyses as this gives a reasonable number of interpretable clusters in both time periods. This was a non-pre-registered researcher decision not based on any a priori rationale. We also experimented with other cluster resolution values (table S1).

**Table S1.**
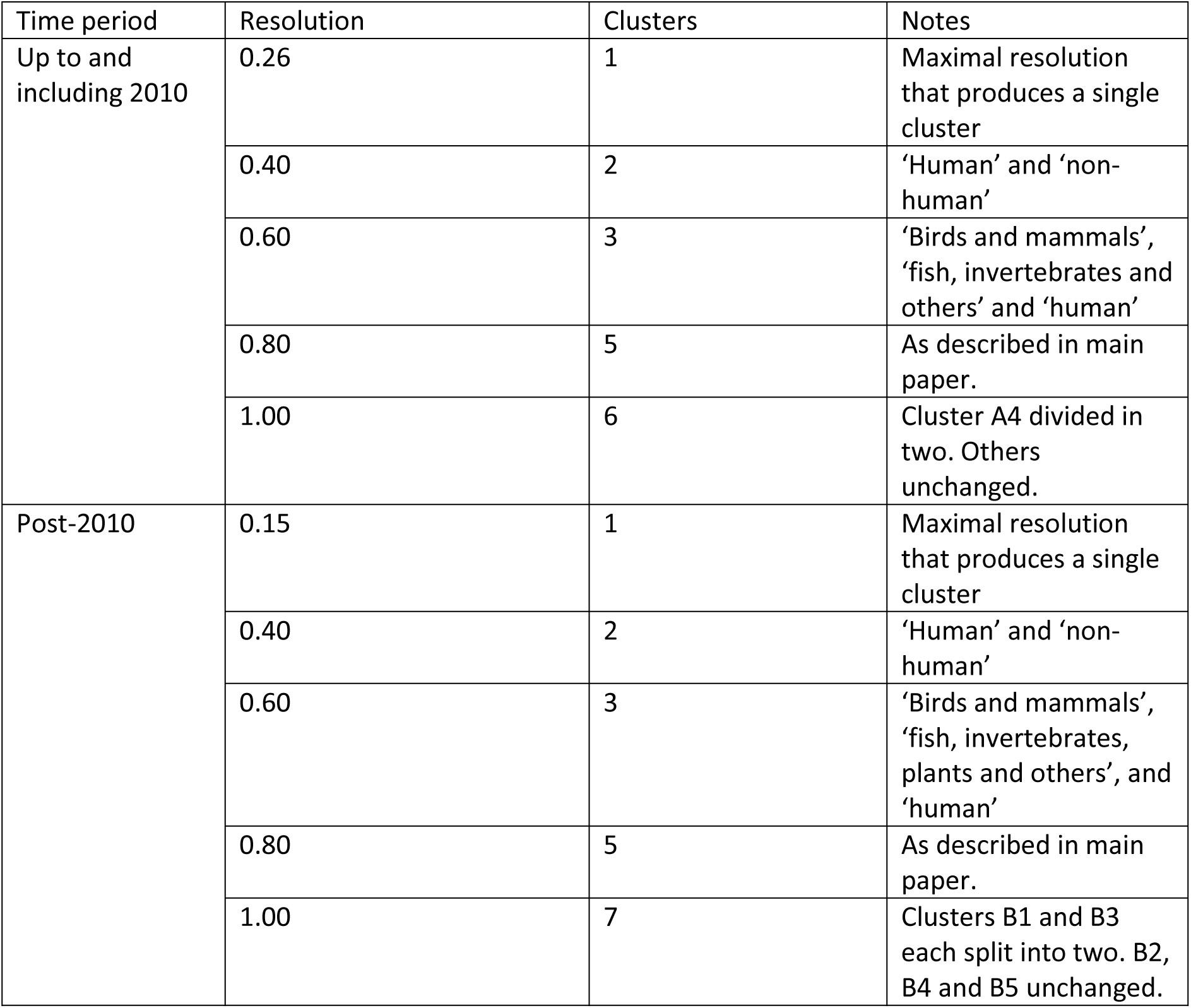
Brief description of cluster structure detected in each time period using different values of the cluster resolution parameter.

The maximum cluster resolution that can be used before distinct clusters were detected was different in the two time periods: 0.26 in the literature up to 2010, but only 0.15 in the more recent period. This, along with the other evidence discussed in the main paper, suggests a greater degree of internal separation in the more recent period. In both time periods, the first cluster division to appear was ‘human’ versus ‘non-human’. At a cluster resolution of 0.60, again in both time periods, the ‘non-human’ cluster split into one focussed on birds and mammals, and another focussed on fish, invertebrates, reptiles, plants, etc. The non-human clusters further divided more finely as the resolution was increased beyond 0.80.

We also examined the maximum cluster resolution that can be used before distinct clusters are detected in the four finer-grained time categories detailed in section 3 of this document. The resolutions were 0.27 (up to and including 2004); 0.25 (2005-2010); 0.18 (2011-14); and 0.16 (2015-8).

